# Co-evolution of dispersal with social behaviour favours social polymorphism

**DOI:** 10.1101/127316

**Authors:** Charles Mullon, Laurent Keller, Laurent Lehmann

## Abstract

Dispersal determines gene flow among groups in a population and so plays a major role in many ecological and evolutionary processes, from biological invasions to species extinctions. Because patterns of gene flow shape kin structure, dispersal is also important to the evolution of social behaviours that influence reproduction and survival within groups. Conversely, dispersal patterns depend on kin structure and social behaviour. Dispersal and social behaviour therefore co-evolve but the nature and consequences of this interplay are not well understood. Here, we model this co-evolution and show that it readily leads to the emergence and maintenance of two broadly-defined social morphs: a sessile, benevolent morph expressed by individuals who tend to increase the fecundity of others within their group relative to their own; and a dispersive, self-serving morph expressed by individuals who tend to increase their own fecundity relative to others’ within their group. This social polymorphism arises as a consequence of a positive linkage between the loci responsible for dispersal and social behaviour, leading to benevolent individuals preferentially interacting with relatives and self-serving individuals with non-relatives. We find that this positive linkage is favoured under a large spectrum of conditions, which suggests that an association between dispersal proclivity and other social traits should be common in nature. In line with this prediction, dispersing individuals across a wide range of organisms have been reported to differ in their social tendencies from non-dispersing individuals.

## Introduction

Dispersal, the movement away from natal habitat to reproduce, is an important step in the life-history of most organisms ^1,2^. At the population level, dispersal patterns shape kin structure which determines whether individuals interact and compete with relatives. This in turn influences the evolution of social behaviour such as helping or aggression ^3,4^. At the same time, dispersal decisions are often influenced by kin and social interactions ^1,5–7^, resulting in the co-evolution among dispersal between groups and social behaviours withingroups ^8–16^. However, the consequences of this co-evolution for within-species behavioural diversity e.g., ^17^ remain elusive. Here, we model the co-evolution between unconditional dispersal and social behaviours, and show that it readily leads to a stable genetic and social polymorphism, whereby individuals who disperse behave differently than non-dispersers.

To model social interactions within groups and dispersal between groups, we assume that the population is structured according to the infinite island model ^5,18,19^, in which individuals belong to local groups and interact socially only with other locals. As a baseline, we assume that groups are of fixed size *N*, that individuals reproduce asexually and then die so that generations do not overlap. An offspring either remains in its natal group (with probability 1 – *d*), or disperses to another randomly chosen one (with probability *d*) and survives dispersal with probability 1 – *c*_d_. Social interactions are modelled with a classical matrix game ^20^: individuals randomly pair up within their group and each independently chooses between two actions denoted B (with probability *z*) and M (with probability 1 – *z*). Depending on the action of each player, each reaps a material payoff that in turn linearly increases its fecundity. Without loss of generality, we assume that when both play M, they obtain no payoff (see Material and Methods M.1 for more details on the game). If one plays B and the other plays M, the B player gets the (direct) benefit *B*_D_ and the M player gets the (indirect) benefit *B*_I_. We assume that *B*_I_ – *B*_D_ *>* 0, which means that an individual who plays B more often that its partner increases its partner’s fecundity relative to its own, and conversely, an individual who plays M more often decreases its partner’s fecundity. We therefore refer to action B as benevolent and M as self-serving. Finally, if they both play B, they each get *B*_D_ *+ B*_I_ – *S*, where *S >* 0 is the antagonistic synergy of benevolence, i.e., *S* captures the degree with which returns diminish with the number of individuals adopting the benevolent action B in a pair.

## Results

First, we study mathematically the co-evolution of the probability *d* of dispersing with the probability *z* of adopting the benevolent action B when they are encoded by two linked loci that experience rare mutations with small quantitative effects ^21^ (M.2 for methods). In agreement with previous results, the population first evolves gradually to converge towards an equilibrium for both traits: dispersal converges to an equilibrium 0 *< d*^*∗*^ *≤* 1 that depends on the cost *c*_d_ of dispersal and group size ^5,22^ (Figure 1), while the probability *z* of adopting the benevolent action B converges to 0 *≤ z*^*∗*^ *= B*_D_/*S ≤* 1 (provided 0 *≤ B*_D_ *≤ S*, S.1.1 for details) ^23^. Once the population has converged to the equilibrium (*d*^*∗*^, *z*^*∗*^) for both dispersal and benevolence, the population either is maintained at this equilibrium by stabilising selection (i.e., the population is uninvadable by any alternative strategy) and remains monomorphic, or undergoes disruptive selection and becomes polymorphic.

**Figure 1:**
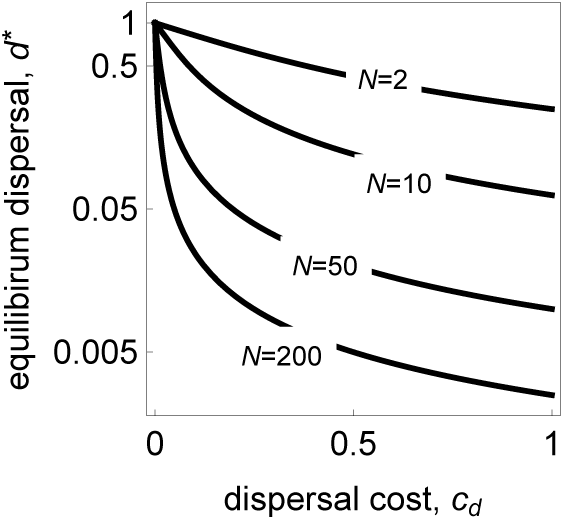
Equilibrium dispersal. Dispersal equilibrium *d*^*∗*^ in terms of the cost of dispersal, *c*_d_, for different group sizes *N* (see Supporting Information S.1.1, eq. S.3 for mathematical expression).

Mathematical analysis reveals that disruptive selection occurs under a wide range of model parameters (Figure 2), and that it leads to the emergence of two morphs: a more benevolent, sessile morph, and a more self-serving, dispersive morph (Supporting Information 1.2-1.3). To understand why selection favours these two morphs, consider an individual that expresses the benevolent, sessile morph. Such an individual tends to preferentially interact with related individuals of the same morph, and so its benevolence is preferentially directed towards relatives. Conversely, an individual from the dispersive morph preferentially interacts with less related individuals, and thus benefits from being self-serving. Polymorphism therefore arises due to the combined effects of dispersal on kin interaction and social behaviour on neighbours’ fitness. In line with this, when only one trait (dispersal or benevolence) evolves and the other is fixed, the population remains monomorphic for all model parameters (Supporting Information 1.2).

**Figure 2:**
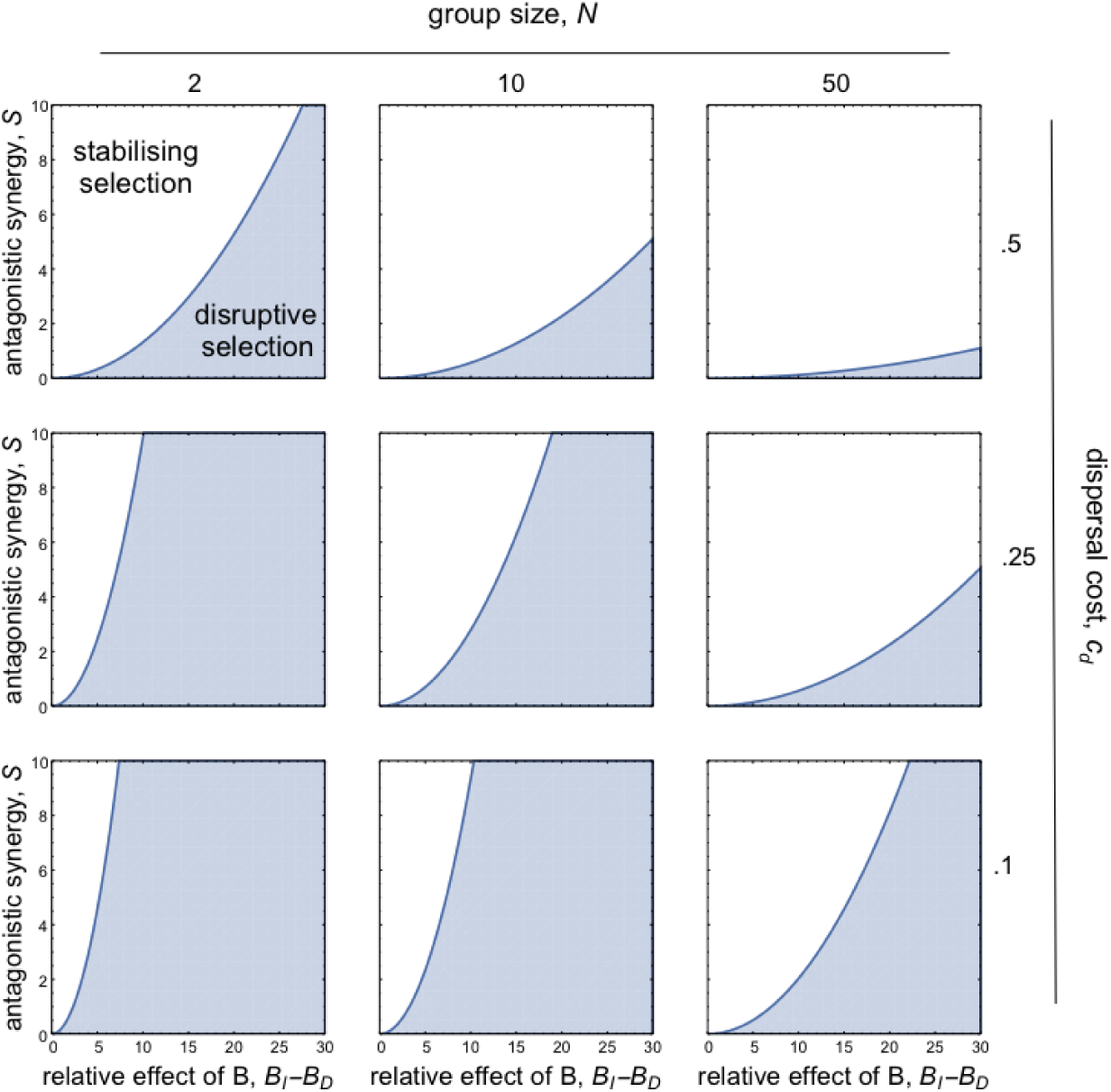
Disruptive selection when dispersal and social behaviour co-evolve. Parameter region under which disruptive selection leads to polymorphism at the equilibrium (*d*^*∗*^, *z*^*∗*^) (shaded region, computed from eq. S.17 in Supporting Information S.1.3, here shown with individual fecundity at the equilibrium set to one). Disruptive selection is therefore favoured when (i) groups are small; (ii) dispersal cost *c*_d_ is low; (iii) antagonistic synergy *S* is weak; and (iv) benevolence has large relative effects (i.e., *B*_I_ – *B*_D_ is large).

To check our mathematical analyses and investigate the long-term effects of disruptive selection, we ran individual-based simulations under conditions that should lead to polymorphism (Supporting Information 3). As predicted, the population first converges to the interior equilibrium for dispersal *d*^*∗*^ and the probability *z*^*∗*^ to adopt benevolent action B, and splits into two morphs, a more benevolent, sessile morph, and a more self-serving, dispersive morph (Figure 3a). Competition among the two morphs then creates a positive feedback that favours more extreme variants. The population eventually stabilises for two highly-differentiated genetic morphs, resulting in a strong association between dispersal and social behaviour (Figure 3b).

**Figure 3:**
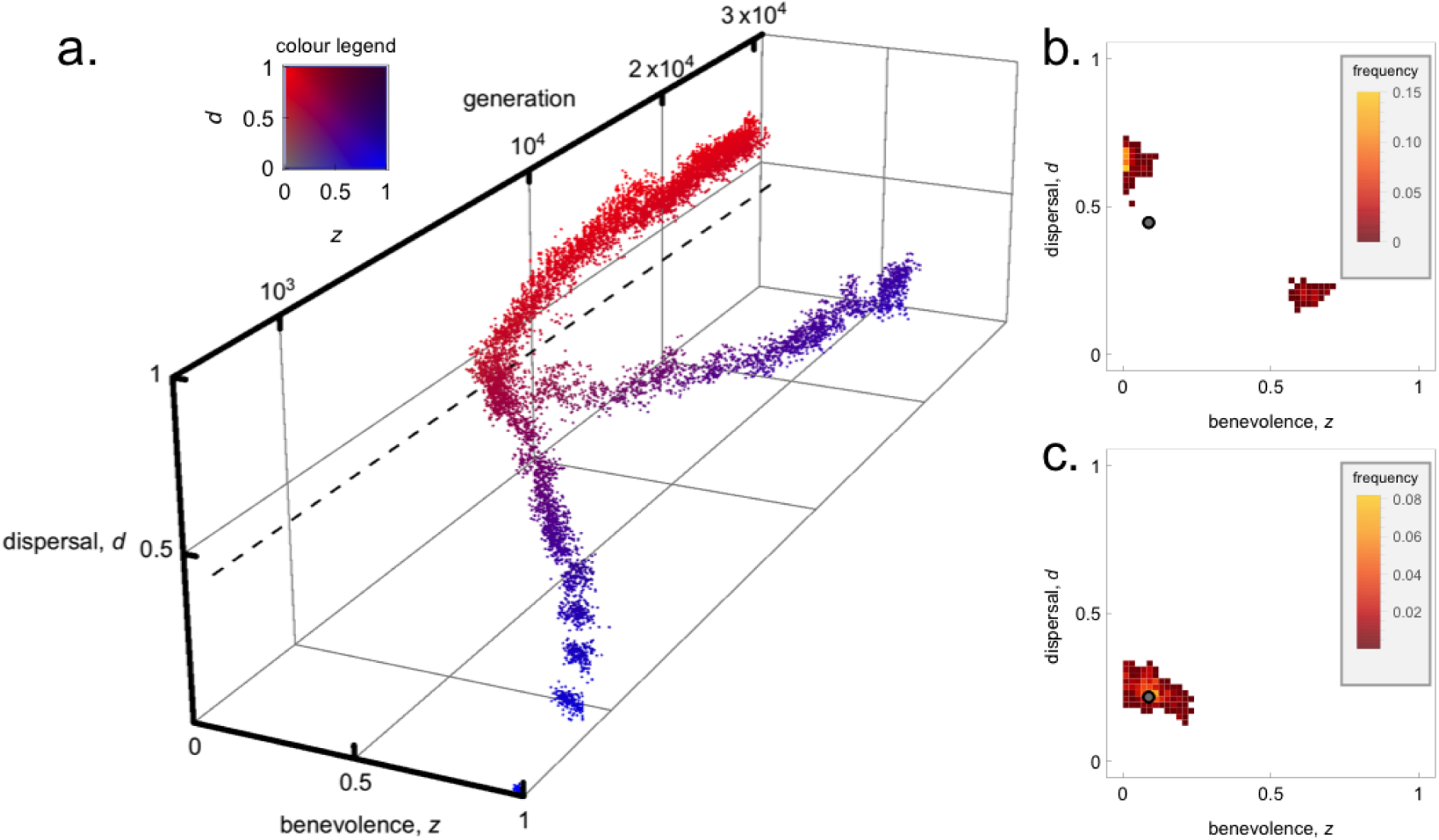
The emergence and maintenance of social morphs when dispersal and social behaviour co-evolve. **a.** Dispersal *d* and benevolence *z* in a simulated population of 10’000 individuals (trait values of 300 individuals randomly sampled every 500 generations, see colour legend for colouring scheme) and predicted interior equilibrium (dashed black line, with *B*_D_ *=* 0.05, *B*_I_ *=* 1.95, *S =* 0.55, *N =* 8, *c*_d_ *=* 0.1, and baseline fecundity set to one). **b.** Phenotypic distribution of whole population at generation 3 *×* 10^4^ (same parameters as a., predicted interior equilibrium shown by grey disc). **c.** Phenotypic distribution of whole population at generation 3 *×* 10^4^ under balancing selection (with *c*_d_ *=* 0.25, other parameters same as a., predicted interior equilibrium shown by grey disc).

To test whether an association between dispersal and social behaviour also emerges when selection is stabilising, we ran simulations under such conditions. As predicted by our analysis, the phenotypic distribution in the population remains centred around the uninvadable equilibrium (Figure 3c). A negative association among dispersal and benevolent behaviour also emerges but is it significantly weaker than when selection is disruptive (compare Figures 3b and c).

Three important assumptions made in the baseline model are that generations do not overlap, that group size is fixed and that the population is structured according to the standard island model. We relaxed the first assumption by performing a mathematical analysis of dispersal and social behaviour co-evolution when a single individual is replaced at each generation in each group. This analysis reveals that polymorphism is also often favoured in this scenario. In fact, compared to our baseline model, polymorphism is favoured for an even greater diversity of payoff variables, which means that a greater diversity of social behaviours may become associated with dispersal when generations overlap (Supporting Information 2). We next relaxed the second and third assumptions by letting group size fluctuate (with regulation through local competition), and by isolating groups by distance (instead of an island model). For both conditions, we compared the outcomes of three simulation experiments: i) we fixed dispersal and let only social behaviour evolve, ii) we fixed social behaviour and let only dispersal evolve, and iii) we allowed both traits to co-evolve. These experiments show that distinct social morphs emerge as a result of disruptive selection, but as under the baseline scenario, this is only true when both traits co-evolve (Supplementary Figures S1-S2).

We also tested the importance of genetic linkage for the emergence of highly differentiated social morphs by studying the effects of various levels of recombination between the loci that control dispersal and social behaviour (Supporting Information 3). This revealed that the emergence of distinct morphs depends on the level of recombination. As recombination increases, maladapted morphs that are benevolent and dispersive or self-serving and sessile increase in frequency (Figure 4a). Beyond a threshold, recombination prevents polymorphism altogether and the population remains monomorphic (Figure 4a). This is because strong recombination breaks the positive genetic linkage among the dispersal and social behaviour loci that is necessary for benevolent individuals to preferentially direct their benevolence towards relatives, and self-serving individuals to compete with non-relatives.

**Figure 4:**
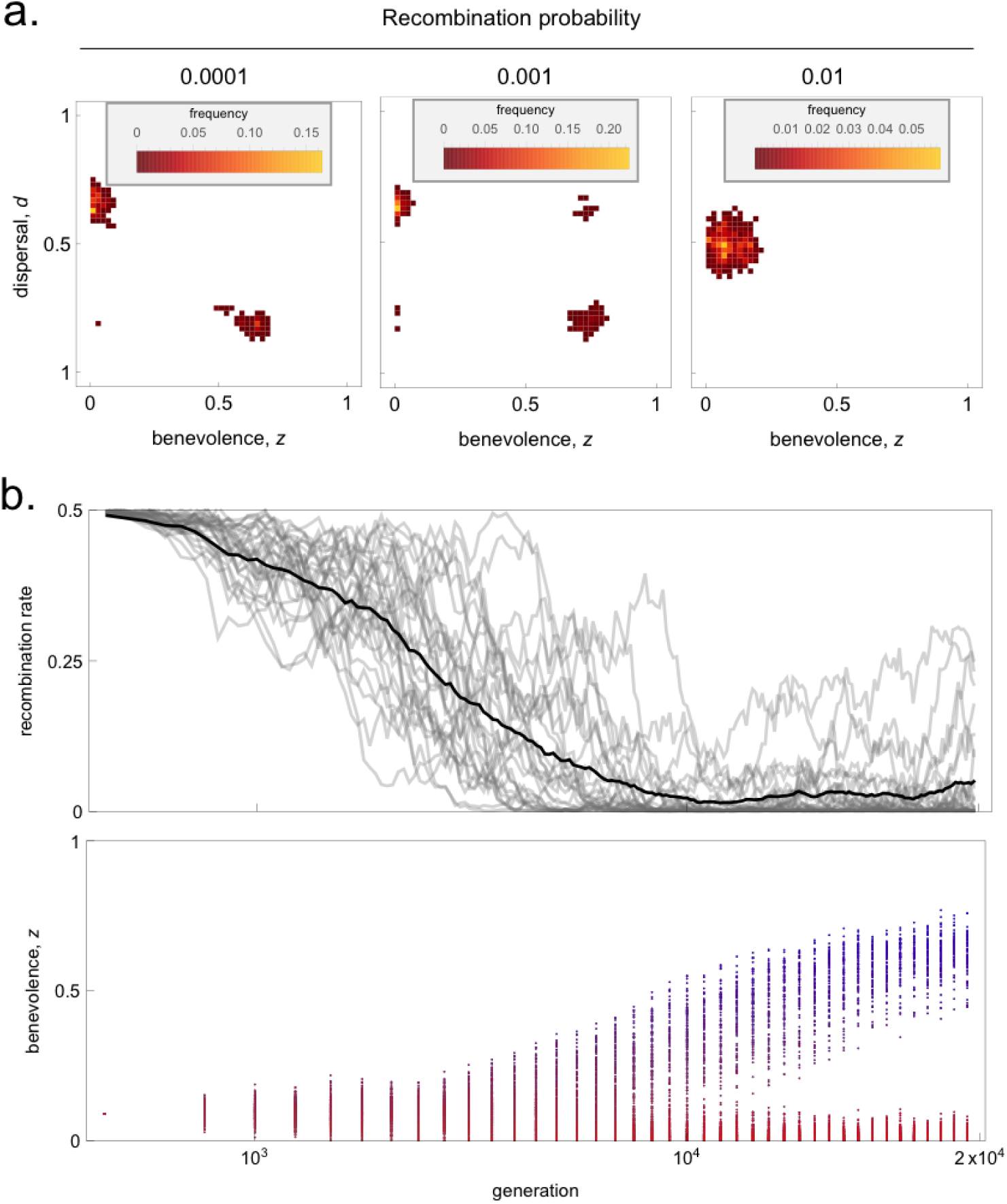
The effect (a) and evolution (b) of genetic linkage between dispersal and social behaviour loci. **A** Phenotypic distribution of whole population at generation 2 *×* 10^4^ when recombination probability is fixed (recombination value shown above graphs, other parameters same as in Fig 3a, Supporting Information 3 for details). **b.** Upper panel: effective recombination rate in the population (number of recombination events/population size) every 100 generations when recombination evolves in a population of 10’000 individuals (for 30 replicates, each replicate is shown by a grey line, average recombination rate over replicates is shown in black). Lower panel: benevolence *z* of 300 individuals randomly sampled every 500 generations across all 30 replicates (coloured according to colour legend in Fig 3). Polymorphism arose in all 30 replicates.

Since disruptive selection promotes an association among dispersal and social behaviour, it should also promote a genetic architecture that makes this association heritable ^24^. We tested this by adding a third locus that controls recombination and let it evolve by introducing two alleles that mutate from one another, one recessive wild-type that codes for a recombination probability of 1/2 and one dominant mutant that stops recombination (Supporting Information 3). Starting with a wild-type population at the predicted equilibrium (*d*^*∗*^, *z*^*∗*^) for dispersal and social behaviour, the mutant allele at the recombination modifier locus eventually invades so that recombination is shut down, which then permits the emergence of distinct morphs (Figure 4b). Disruptive selection therefore leads to the genetic integration of dispersal and social behaviours to form a "supergene" ^25^, which allows benevolent individuals to preferentially interact with relatives, and self-serving individuals with non-relatives. This type of kin association through genetic and spatial assortment may constitute a first step towards conditional dispersal ^10^ or conditional social behaviours ^8,11,26^, which allow individuals to fine tune their behaviours towards relatives.

Finally, we studied the population genetic signatures associated with the emergence of this social polymorphism. We first calculated the degree of genetic differentiation (*F*_ST_) among morphs at the locus responsible for social behaviour (the "selected locus") and at a neutral, unlinked locus. Genetic differentiation among morphs at the selected locus is much greater than at a neutral locus (Figure 5), which is unsurprising since social behaviour is genetically determined. Second, we looked at the degree of genetic differentiation among groups within each social morph, and found that differentiation among groups is greater within the benevolent morph than the self-serving one at the selected locus (Figure 5). This pattern arises because the benevolent morph has lower dispersal tendencies than the self-serving one. Interestingly, differentiation among groups is also greater within the benevolent morph at the neutral locus (Figure 5). The effect of dispersal differences between the social morphs therefore extends beyond the selected locus and creates detectable patterns of genetic differentiation at neutral unlinked loci.

**Figure 5:**
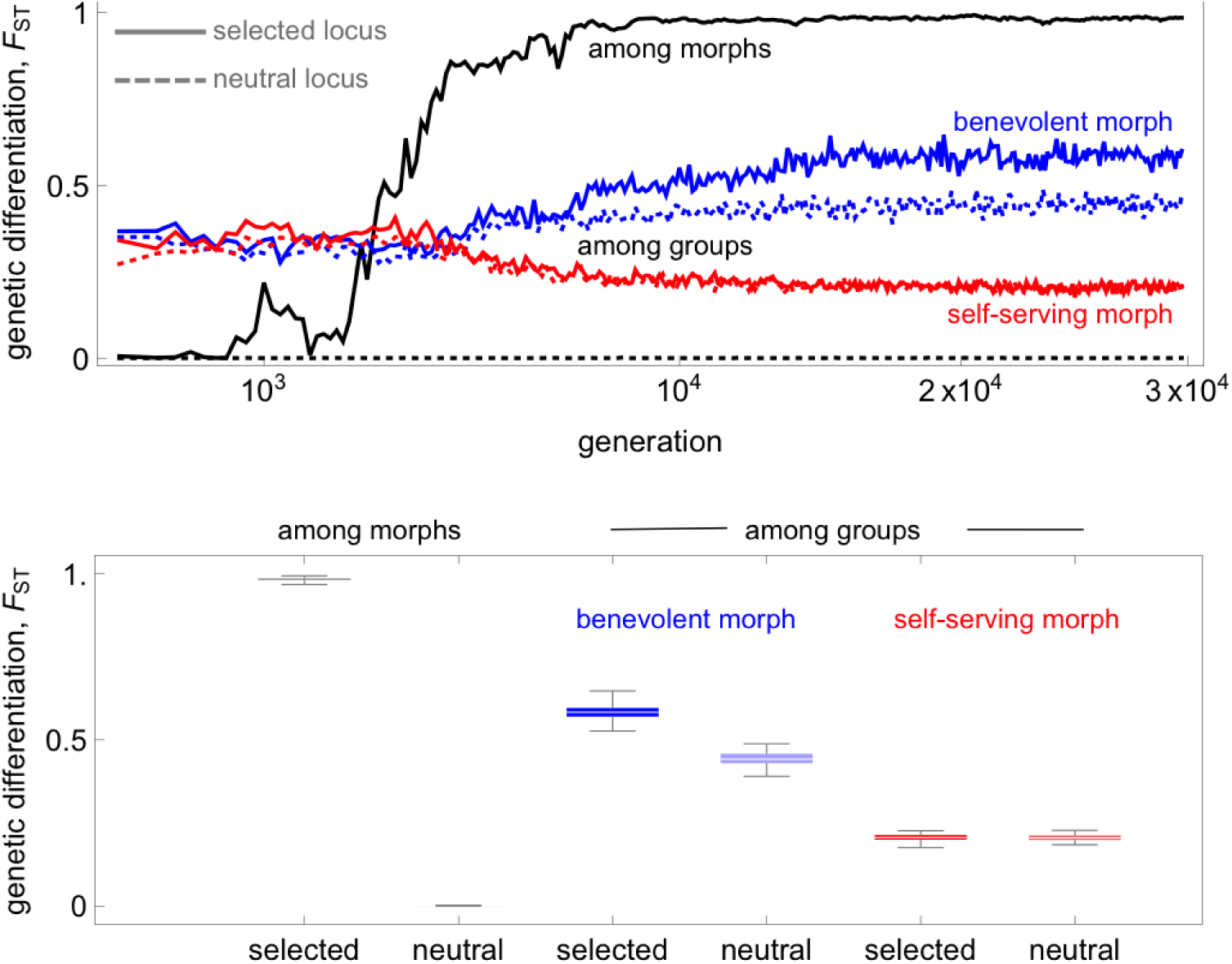
Patterns of genetic differentiation. Upper panel. *F*_ST_ values calculated every 100 generations for a population initially monomorphic for its predicted equilibrium that becomes polymorphic when the loci for dispersal and social behaviours are genetically linked (see Supplementary Information S.3 for information on *F*_ST_ calculations, simulation parameters same as in Fig 3a). Black: genetic differentiation among morphs at the selected locus (full line) and at an unlinked neutral locus (dashed line). Blue: genetic differentiation among groups within the benevolent morph at the selected locus (full line) and at an unlinked neutral locus (dashed line). Red: genetic differentiation among groups within the self-serving morph at the selected locus (full line) and at an unlinked neutral locus (dashed line). **Lower panel:** summary of the *F*_ST_ values at equilibrium (be-tween generations 1.5 *×* 10^4^ and 3 *×* 10^4^) among morphs (black) and among groups within benevolent (blue) and self-serving (red) individuals at the selected (darker shade) and an unlinked neutral (lighter shade) locus.

## Discussion

These analyses reveal that the co-evolution of dispersal and social behaviour favours the emergence of a social polymorphism and dispersal syndrome. Clear predictions can be extrapolated from our results. The first is that phenotypic associations between dispersal and social behaviour should be common. This prediction finds echo in multiple organisms ranging from protozoa to primates where an intraspecific association between dispersal and social behaviour have been found (Table 1, see also ^10,27–29^ for reviews). For example, in the ciliate *Tetrahymena thermophila*, laboratory studies have shown that strains that are more cooperative disperse at a lower rate than less cooperative strains at intermediate population densities ^30^. Similarly, in wild populations of prairie voles *Microtus pennsylvanicus*, individuals that disperse tend to be more aggressive than those who do not ^31^.

**Table 1:** Species where a phenotypic association between dispersal and social behaviour has been described.

| Social behaviour | Association with dispersal | Species |
| --- | --- | --- |
| Aggressiveness,<br>"Self-servingness" | Positive | *Sialia mexicana*<sup>59,60</sup>, *Clethrionomys rufocanus*<sup>61</sup>, *Microtus pennsylvanicus*<sup>31</sup>, *Myodes glareolus*<sup>62</sup>, *Rhabdomys pumilio*<sup>63</sup>, *Neolamprologus pulcher*<sup>64</sup>, *Mus musculus domesticus*<sup>65</sup>, *Macaca mulatta*<sup>66–68</sup>, *Drosophila melanogaster*<sup>35</sup> |
| Pro-social,<br>Helping,<br>Cooperation,<br>"Benevolence" | Negative | *Picoides borealis*<sup>69</sup>, *Lacerta vivipara*<sup>70</sup>, *Tetrahymena thermophila*<sup>30</sup>, *Uta stansburiana*<sup>71</sup>, *Heterocephalm glaber*<sup>72</sup>, *Apus melba*<sup>36</sup> |

Our prediction that benevolence and dispersal propensity should be associated at the phenotypic level aligns with the predictions stemming from models of conditional behaviours ^8,10,11^. When individuals can condition their social behaviour on whether they have dispersed or not, selection favours increased benevolence in dispersers and self-servingness in non-dispersers ^11^, also creating a negative phenotypic association between benevolence and dispersal behaviours. Although studies on conditional behaviours ^8,10,11^ have not tested whether genetic polymorphism would also emerge, such an association is unlikely because conditional behaviour already allows to behave more benevolently towards relatives. The evolution of conditional behaviour thus results in a phenotypic but not a genetic association among dispersal and social behaviours.

By contrast, the social polymorphism we report here is underlain by a genetic association among dispersal and social behaviours. Previous models of the co-evolution of unconditional dispersal and cooperation have also found that genetic polymorphism can emerge in asexual haploids ^14–16^. But these models assume that cooperation has non-linear fitness effects such that polymorphism can arise when cooperation is evolving but the rate of dispersal is fixed. As a result, the importance of dispersal and social behaviours co-evolution for polymorphism is unclear from these models. By contrast, in our model, a polymorphism only emerges when both traits co-evolve.

Importantly, our analyses suggest that selection also leads to the genetic integration of dispersal and social behaviours into a single Mendelian genetic element. This brings us to our second prediction: dispersal and social behaviour should be genetically associated when polymorphism is present. Data to test this prediction are scarce because few studies combine dispersal, behavioural and heritability assays. One notable exception is found in western bluebirds *Sialia mexicana*, for which a multi-generational pedigree analysis revealed that dispersal and social behaviour are genetically associated such that dispersive males are more likely to produce aggressive offspring and non-dispersive males, non-aggressive offspring ^32^. A similar pattern of positive genetic association among dispersal and aggressiveness has been observed in two species of wild house mice *Mus musculus musculus* ^33^ and *Mus domesticus* ^34^. Lines of *Drosophila melanogaster* that have been selected for greater dispersal propensity also exhibit elevated aggressiveness ^35^, which shows that genetic correlations among dispersal and aggressiveness are also present in this species. Conversely, in the colony-breeding Alpine swift *Apus melba*, dispersal and cooperative defence against human intrusion are negatively associated both at the phenotypic and genetic level ^36^.

Our prediction of a genetic association among dispersal and social behaviour could be tested further in so-cially polymorphic species, such as social spiders or halictine bees. Between *Anelosimus* spider species, natal dispersal tends to be negatively associated with social living and behaviour ^37^. It would be interesting to test whether this association also occurs within species. In particular in the social spider *Anelosimus studiosus* an aggressive morph co-exists with a docile one ^38^, and evidence already suggests that expressing the aggressive morph is heritable ^39^ and associated with other social behaviours ^40^, but an association between dispersal and aggressiveness has not yet been studied. Direct evidence for such an association would require dispersal and heritability assays that can be challenging. Alternatively, indirect evidence could be provided by patterns of genetic differentiation (*F*_ST_) among groups. In particular, our model suggests that a genetic association between dispersal and aggressiveness should lead to greater between-group genetic differentiation (at both selected and neutral loci) for the aggressive morph than for the docile morph (Figure 5).

Eusocial species, who typically exhibit rich and variable patterns of dispersal and social behaviours, also provide a good model to test our prediction that dispersal and social behaviour should be genetically associated. Many ant species show a dispersal syndrome that associates dispersal with social organisation. Queens either disperse far away from their natal nest and form single-queen (monogyne) colonies, or disperse short distances and form multiple-queen (polygyne) colonies ^41^. These two morphs are frequently found in the same population and show little genetic differentiation, suggesting extensive gene flow among them ^41–44^. In line with the predictions of our model, individuals from monogyne colonies exhibit high intra-specific aggression towards non-nestmates while individuals from polygyne colonies are much less aggressive ^41^. The genetic underpinning of social organisation has been uncovered in two ant species, *Solenopsis invicta* ^43^ and *Formica selysi* ^44^. Remarkably, in both cases, the social polymorphism and dispersal syndrome is controlled by a large non-recombining region that has independently arisen in each species, which suggests that integration of dispersal and social behaviour into a supergene can readily occur in nature.

Our simple model of course cannot explain all associations among dispersal and social behaviour which can be influenced by many other factors (e.g., in species with a social hierarchy, such as meerkats ^45^, it may be beneficial to be more submissive when dispersing into a foreign group, so that we may expect benevolent behaviours to be positively associated with dispersal, see also ^27,29^). Yet, the selection that associates dispersal and social behaviour in our model will influence evolution under most ecological settings because it only depends on kin structure which, due to limited dispersal and the spatial scale of social interactions, is ubiquitous in nature ^46^. While current data support the notion that individuals who disperse behave towards conspecifics in a way that is different to non-dispersers, further pedigree and genomic analyses will provide a better picture of how associations among dispersal and social behaviour are genetically constructed.

## Materials and Methods

### M.1 Matrix game

We use a pairwise symmetric matrix game ^20^ to model social interactions within groups. Without loss of generality, we assume that the game is described by the following payoff matrix

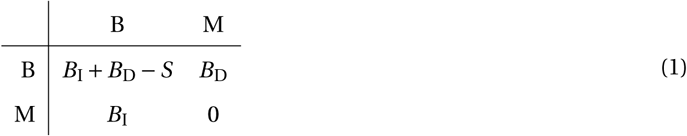

whose entries give the payoff to a focal row player (i.e., when both play B, the focal obtains *B*_I_ *+ B*_D_ – *S*; when the focal plays B and the partner plays M it obtains *B*_D_; in the reverse situation, the focal obtains *B*_I_; and when both partners play M, the focal obtains 0).

The payoff matrix (eq. 1) entails that the average payoff to a focal player who adopts action B with probability *z*_1_ against a partner who adopts this action with probability *z*_2_ is

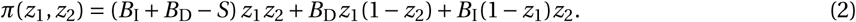

We assume that the payoff that an individual receives increases its fecundity linearly, in which case the fecundity of the partner, relative to the fecundity of the focal player, can be written as

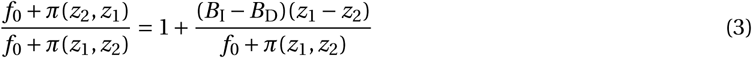

where *f*_0_ is a baseline fecundity that ensures that fecundity is positive. Eq. (3) shows that when *B*_I_ – *B*_D_ *>* 0 and the focal is more likely to express B than its partner (*z*_1_ – *z*_2_ *>* 0), this results in an increase of the partner’s fecundity relative to the focal’s. Conversely, when the focal is less likely to express B than its partner (*z*_1_ – *z*_2_ *<* 0), this results in a decrease of its partner’s fecundity relative to its own. The quantity *B*_I_ – *B*_D_ can therefore be thought of as the relative fecundity effect of action B.

We assume throughout that *B*_I_ – *B*_D_ *>* 0 so that expressing action B increases the fecundity of its recipients (for all *z*_^2^_ *<* 1) relative to its actor, and we therefore call behaviour B "benevolent". Conversely, expressing M increases the fecundity of its actor relative to its recipient (for all *z*_^2^_ *>* 0), and we call behaviour M "self-serving". The condition that *B*_I_ – *B*_D_ *>* 0 includes well-known social dilemma games. For instance, when *B*_I_ *> B*_I_ *+ B*_D_ – *S > B*_D_ *>* 0, B behaviour is "Dove" in the Hawk-Dove game, or "Cooperate" in the Snow-Drift (or Volunteers’ dilemma) game. When *B*_I_ *> B*_I_ *+ B*_D_ – *S >* 0 *> B*_D_, B is "Cooperate" in the Prisoner’s dilemma game. Behaviour B therefore generally encompass cooperative and altruistic behaviours but not necessarily. Whether behaviour B is cooperative or altruistic *sensu* evolutionary biology depends on its fitness effects ^19,47^, which themselves depend on population structure and life-cycle.

## M.2 Evolutionary invasion analysis in the island model

### Invasion and the average mutant growth rate

In the infinite island model, the fate of a mutation that codes for a rare mutant phenotype **x**_m_ *=* (*z*_m_, *d*_m_) when the resident population has phenotype **x** *=* (*z*, *d*) can be deduced from the geometric growth rate *W* (**x**_m_, **x**) of that mutation ^48,49^, which is the time-averaged mean cumulative growth over different replicates or sample paths of the invasion dynamics. If the geometric growth rate is less or equal to one (*W* (**x**_m_, **x**) *≤* 1), then the mutation will eventually go extinct in the population, otherwise it may persist indefinitely ^16^. When the mutant and residents only differ by a small amount (‖**x**_m_ - **x** *≪* 1‖), the growth rate can be approximated by Taylor expanding *W* (**x**_m_, **x**) close to resident phenotype **x**,

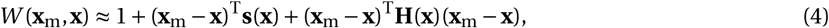

where **s**(**x**) is a 2 *×* 1 vector and **H**(**x**) is a 2 *×* 2 matrix that respectively give the first- and second-order effects of selection ^16,49^, which can be used to infer on the adaptive dynamics of both traits. We detail **s**(**x**) and **H**(**x**) below.

### Directional selection in the infinite island model

When mutations are rare with weak phenotypic effects, the population first evolves under directional selection whereby selected new mutations rapidly sweep the population before a new mutation arises, so that the population “jumps" from one monomorphic state to another ^21^. The direction of evolution under directional selection is indicated by the selection gradient vector

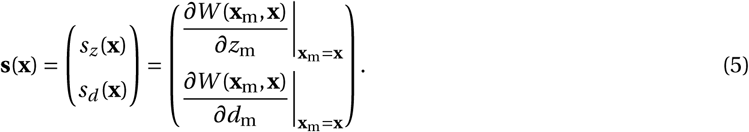

In the infinite island model, the selection gradient on trait *u ∈* {*z*, *d*}, which captures the directional coefficient of selection on trait *u*, has been shown ^e.g., eq. 12 of ref. 16^ to be equal to

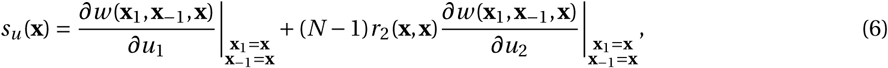

where *w* (**x**_^1^_, **x**_–1_, **x**) is the individual fitness of a focal individual that we arbitrarily label as individual "1" (i.e., the expected number of adult offspring produced by individual "1"), when it has phenotype **x**_^1^_ *=* (*z*_1_, *d*_1_), his *N -* 1 neighbours have phenotypes **x**_–1_ *=* (**x**_2_,…, **x**_*N*_), and the resident has phenotype **x**; and *r*_*l*_ (**x**_m_, **x**) is defined as the probability that *l -* 1 randomly drawn (without replacement) neighbours of a mutant are also mutants (i.e., that they all belong to the same lineage). In a monomorphic population (so that **x**_m_ *=* **x**), *r*_*l*_ (**x**, **x**) reduces to the probability of sampling *l* individuals without replacement whose lineages are identical-by-descent, which is the standard *l* ^th^-order measure of relatedness for the island model ^50^. The selection gradient (eq. 6) is therefore the sum of the direct fitness effects of trait *u* and the pairwise relatedness (*r*_^2^_(**x**, **x**)) weighted indirect fitness effects of trait *u* (note: **x**_–1_ *=* **x** means **x**_2_ *=* **x**_3_ *= =* **x**_*N*_ *=* **x**) ^16,19^.

In two-dimensional phenotypic space, **s**(**x**) points towards the direction of directional selection close to the resident, so adaptive dynamics will first settle for an equilibrium

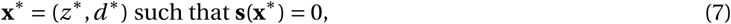

when the equilibrium is an attractor of selection. The condition for the equilibrium **x**^*∗*^ to be a local attractor depends on whether the two traits are genetically correlated. When traits are not genetically correlated (so that mutations have independent effects on both traits) and mutations affect only one trait at a time (no pleiotropy), the equilibrium is a local attractor of the evolutionary dynamic if the Jacobian matrix,

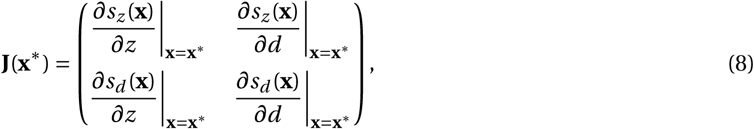

has all eigenvalues with negative real parts. More generally, in the presence of pleiotropy and/or genetic correlations among traits (so that mutations have correlated effects on both traits), the equilibrium **x**^*∗*^ is a local attractor if the symmetric part of the Jacobian matrix **J**(**x**^*∗*^) (eq. 8),

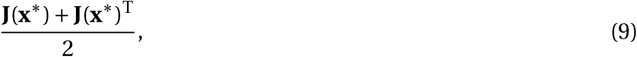

is negative-definite (i.e., if it has only negative eigenvalues), and such an equilibrium is referred to as (strongly) convergence stable ^51,52^. Note that if the symmetric part of the Jacobian matrix **J**(**x**^*∗*^) (eq. 9) is negative-definite, then **J**(**x**^*∗*^) has eigenvalues with negative real parts. For polymorphism to emerge when mutations have weak effects, it is necessary that the population is first at a convergence stable equilibrium ^53^.

### Stabilising/disruptive selection in the infinite island model

Once the population is at an equilibrium **x**^*∗*^ that is convergence stable, the leading eigenvalue *λ*(**x**^*∗*^) of the Hessian matrix,

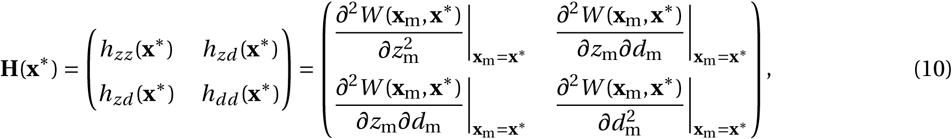

tells us whether selection is stabilising (when *λ*(**x**^*∗*^) *≤* 0 and all mutations close to the resident vanish) or disruptive (when *λ*(**x**^*∗*^) *>* 0 and polymorphism emerges). Note that the Hessian necessarily has real eigenvalues because it is symmetric with real entries.

In the infinite island model, it has been shown ^eq. 13 of ref. 16^ that the *h*_*uv*_ (**x**^*∗*^) entry of the Hessian for *u ∈* {*z*, *d*} and *v ∈* {*z*, *d*}, which is the quadratic coefficient of selection on traits *u* and *v*, can be decomposed as

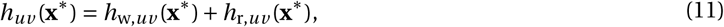

where

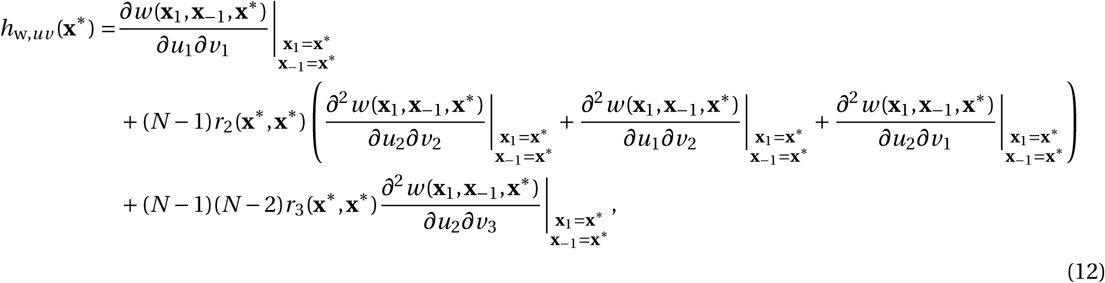

captures the second-order fitness effects when the relatedness among mutants is the same as among residents (since *r*_2_(**x**^*∗*^, **x**^*∗*^) and *r*_3_(**x**^*∗*^, **x**^*∗*^) are evaluated when the population is monomorphic for the resident at equilibrium), and where

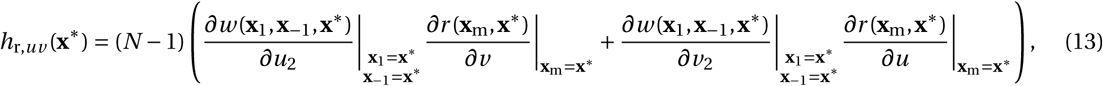

depends on the effects that the traits have on pairwise relatedness (i.e., on *∂r* (**x**_m_, **x**^*∗*^)/*∂v* and *∂r* (**x**_m_, **x**^*∗*^)/*∂u*). In order to evaluate this effect, it is first necessary to decompose individual fitness *w* (**x**_1_, **x** -_1_, **x**) as

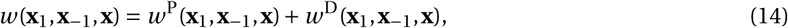

where *w* ^P^(**x**_1_, **x**-_1_, **x**) is the expected number of offspring of the focal individual that remain in their natal group (i.e., its expected number of philopatric offspring), and *w* ^D^(**x**_1_, **x** -_1_, **x**), is its number of offspring that disperse. Then, for the models considered here, previous works have shown that the effect of trait *v ∈* {*z*, *d*} on relatedness can be expressed as

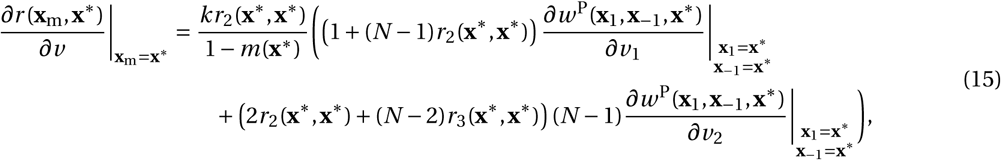

where *k* is a constant that depends on the life-cycle (*k =* 2 for the baseline Wright-Fisher model ^eq. 18 of ref. 54, eq. 28 of ref. 55^, *k = N* when generations overlap under the Moran model ^eq. 14 of ref. 16^) and *m*(**x**^*∗*^) is the neutral backward probability of dispersal (i.e., the probability that a breeding spot is filled by an immigrant in a population monomorphic for the resident).

The quadratic coefficient of selection on a single trait (*h*_*zz*_ (**x**^*∗*^) and *h*_*dd*_ (**x**^*∗*^)) tell us about selection on that trait when it is evolving in isolation from the other. For instance, when *h*_*zz*_ (**x**^*∗*^) *≤* 0, selection on *z* is stabilising, but when *h*_*zz*_ (**x**^*∗*^) *>* 0 selection is disruptive and *z* will diversify whether or not dispersal is also evolving. Meanwhile, the quadratic coefficient of selection on *z* and *d*, *h*_*zd*_ (**x**^*∗*^), captures the types of associations or correlations among *z* and *d* that are favoured by selection. It is therefore referred to as the correlational coefficient of selection ^56^. When *h*_*zd*_ (**x**^*∗*^) is positive, selection favours a positive correlation among both traits close to the resident and conversely, when it is negative, selection favours a negative correlation. It follows from standard linear algebra results ^57^, that if

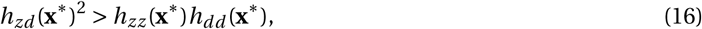

then the leading eigenvalue of **H**(**x**^*∗*^) is positive (*λ*(**x**^*∗*^) *>* 0), which in biological terms means that if the correlational coefficient of selection is strong relative to the quadratic coefficient of selection on both traits, it causes selection to be disruptive and hence, polymorphism.

### Fitness and genetic structure for baseline model

Here, we give the necessary components to perform an invasion analysis for dispersal and benevolence under the baseline model in which generations do not overlap (i.e., Wright-Fisher life cycle). First, note that the fecundity of the focal individual "1" is

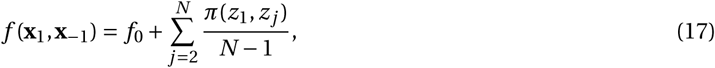

where the payoff function *π* is given in eq. (2). Then, under the baseline model, the expected number of philopatric of individual "1" is

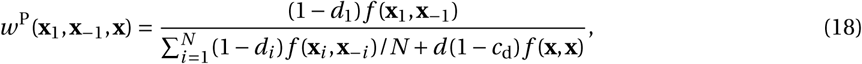

and the overall fitness of individual "1" is

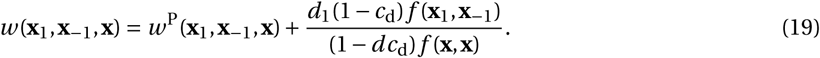

In order to express pairwise and three-way relatedness (eqs. 6–15), we first give the neutral backward probability of dispersal (eq. 15):

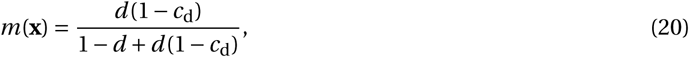

which is the ratio of the number immigrant offspring to the total number of offspring in a group. Pairwise and three-way relatedness are found using *m*(**x**) and standard identity-by-descent arguments ^19,58, eqs. 13 & 23 of ref. 55^, yielding

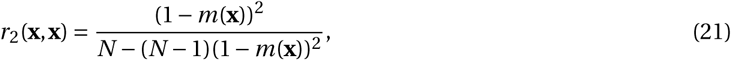

and

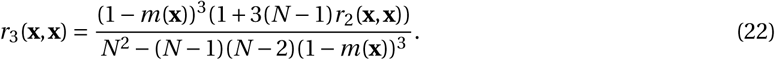

This is all that is necessary to infer on the adaptive dynamics of dispersal and benevolence using the selection gradient and Hessian matrix (eqs. 6–15). Analysis can be found in Supporting Information 1.

### Fitness and genetic structure for baseline model when generations overlap

To incorporate generational overlap, we assume that after reproduction, a random individual in each group dies and that offspring then compete for the single open breeding spot left vacant in each group at each generation (as in a birth-death local Moran process). In this case, fecundity and neutral backward dispersal are as above (eqs. 17 and 20). But philopatric fitness and total individual fitness are now respectively given by

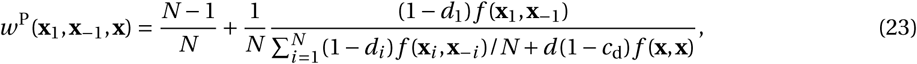

and,

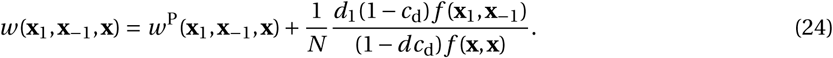

e.g., 16. Meanwhile, pairwise and three-way relatedness ^Table 1 of ref. 16^ are given by

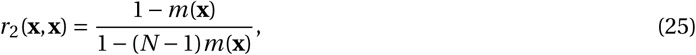

and

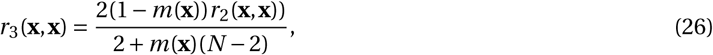

which completes the necessary expressions to derive the adaptive dynamics of dispersal and benevolence when generations overlap. Analysis can be found in Supporting Information 2.

## Supplementary Figures

**Supplementary figure S1:**
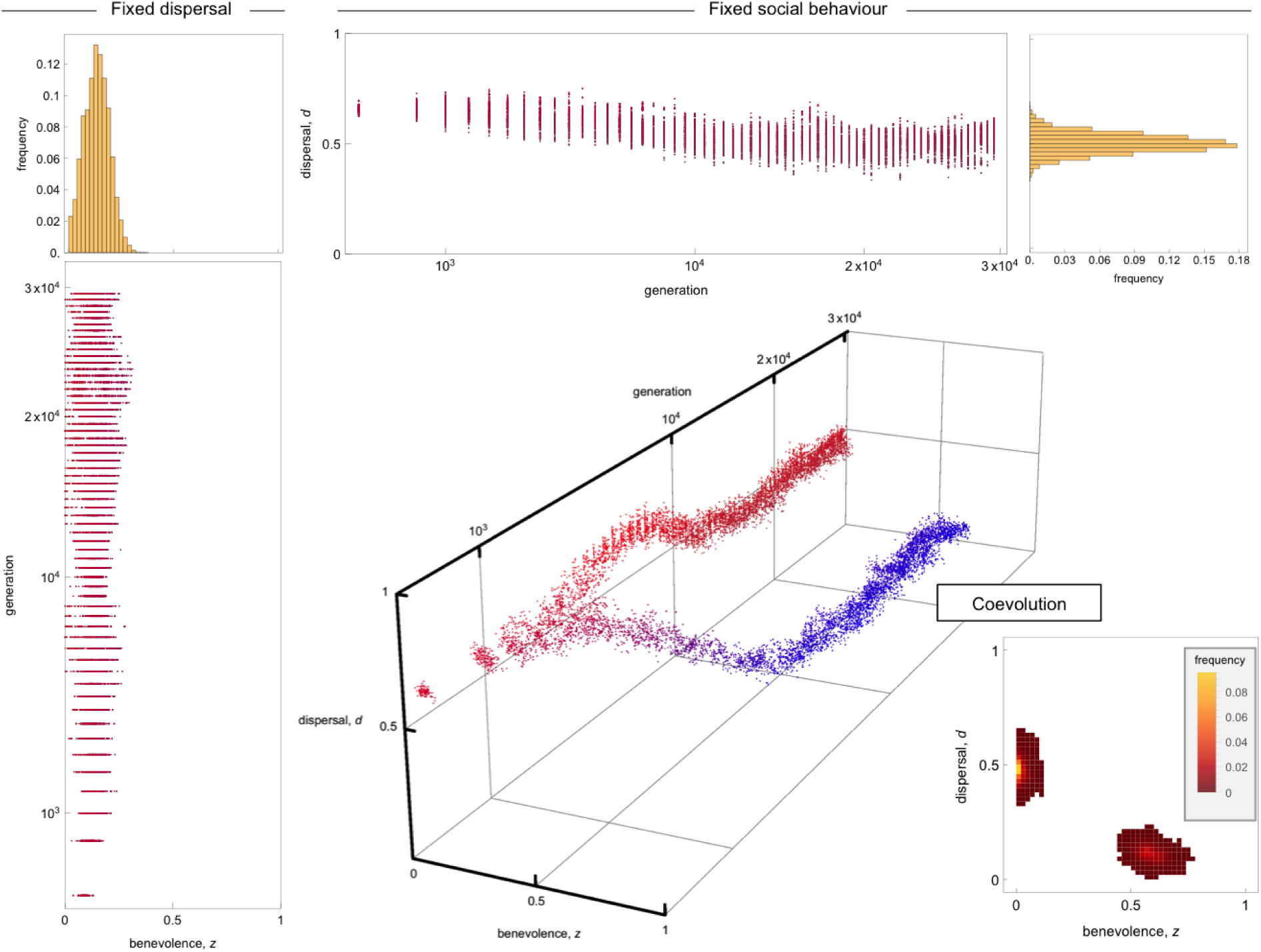
Evolution of dispersal and social behaviour under group size fluctuations. Group size fluctuations were implemented by assuming that before dying, each adult individual produced a Poisson distributed number of offspring, with mean given by eq. (17) multiplied by a factor of 5 to avoid extinction. Each dispersing offspring independently survives dispersal with probability 1 – *c*_d_ (*c*_d_ *=* 0.05 here). After dispersal, offspring compete locally to survive to adulthood: each survive with a probability 1/(1 *+ ηN*_off_), where *N*_off_ is the number of offspring coming into competition (*η =* 0.04, other parameters same as in Fig 3a). Left column: evolution of the probability *z* of adopting benevolent behaviour when dispersal is fixed at *d =* 0.65 and equilibrium distribution of *z* (between generations 1.5 *×* 10^4^ and 3 *×* 10^4^). Top row: evolution of the dispersal probability *d* when *z* is fixed at *z =* 0.1 and equilibrium distribution of *d*. Bottom right: co-evolution of *z* and *d* and equilibrium distribution.

**Supplementary figure S2:**
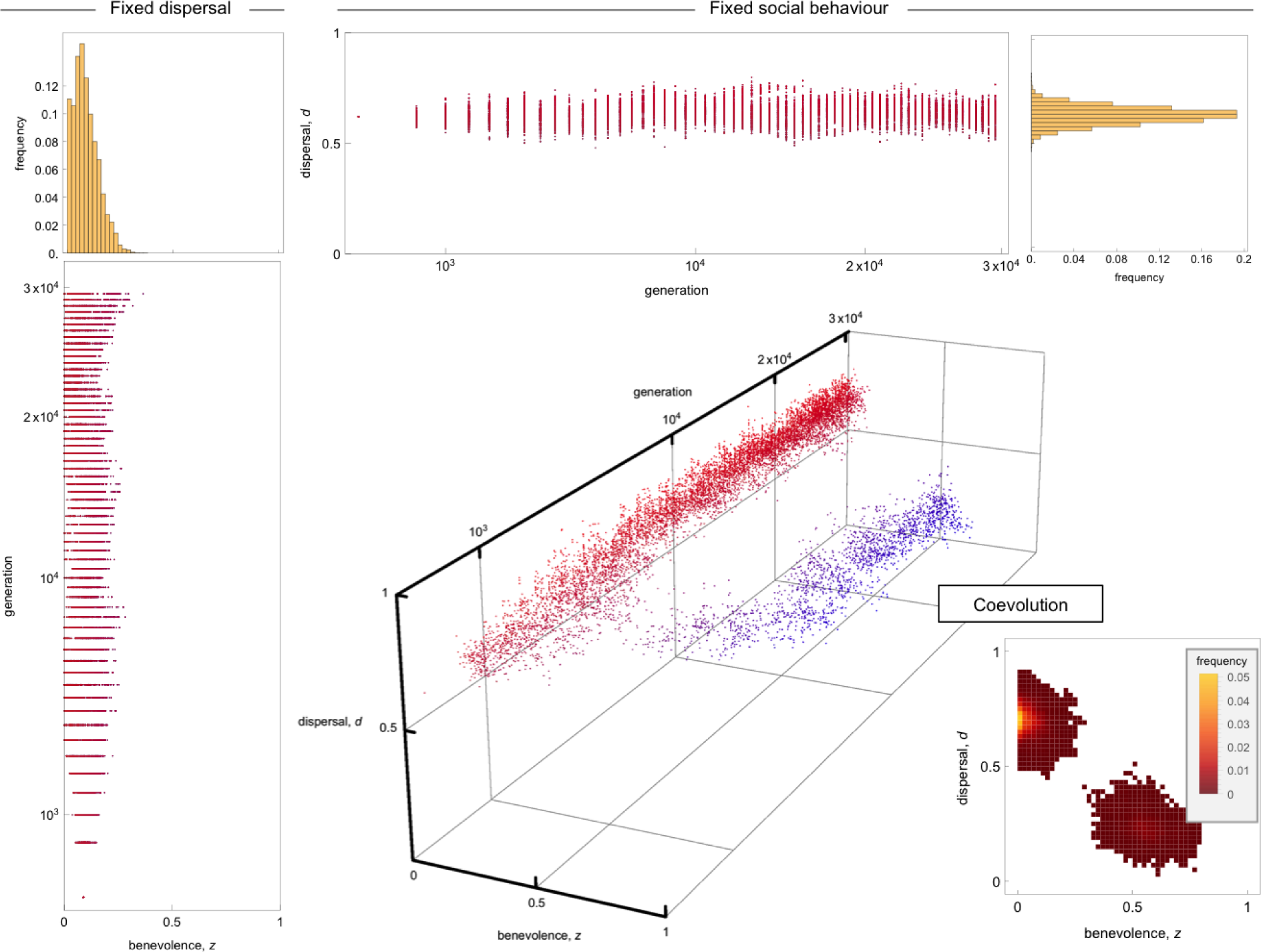
Evolution of dispersal and social behaviour under isolation-by-distance. To incorporate isolation-by-distance, we assumed that social groups are arranged on a 2-D lattice (size: 35 by 35 groups), shaped as a taurus to avoid boundary effects. A dispersing offspring lands to one of the nearest neighbouring group with equal probability (dispersal distance is set to one). The rest of the life-cycle is the same as in our baseline scenario (*c*_d_ *=* 0.05, other parameters same as in Fig 3a). Left column: evolution of the probability *z* of adopting benevolent behaviour when dispersal is fixed at *d =* 0.65 and equilibrium distribution of *z* (between generations 1.5 *×* 10^4^ and 3 *×* 10^4^). Top row: evolution of the dispersal probability *d* when *z* is fixed at *z =* 0.1 and equilibrium distribution of *d*. Bottom right: co-evolution of *z* and *d* and equilibrium distribution.

**Supplementary figure S3:**
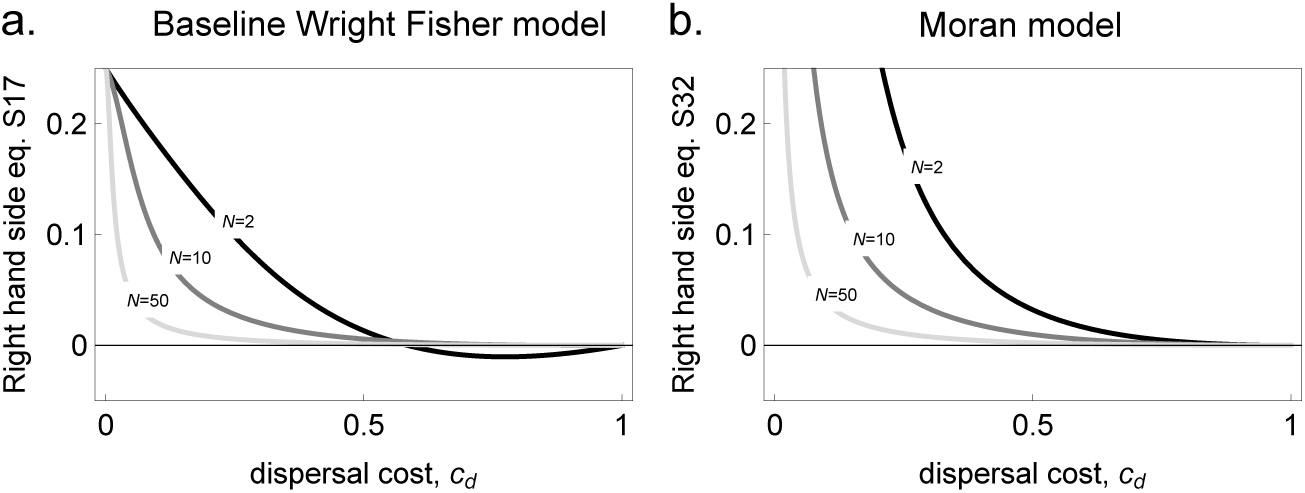
Necessary conditions for disruptive selection in (a) the baseline model and (b) the Moran model. For disruptive selection to occur, it is necessary that the quantity *S f* (**x**^*∗*^, **x**^*∗*^)/(*B*_I_ – *B*_D_)^2^ lies between zero and the plotted curve, shown as a function of dispersal cost and for different group sizes (see eqs. S.17 and S.32). In both models, low dispersal cost and small group sizes thus favour disruptive selection. Because the area between zero and the plotted curve for all dispersal costs and group sizes is larger under the Moran model (compare a and b), disruptive selection is more likely to occur in this model.

## Supporting Information

### S.1 Adaptive dynamics for the baseline Wright-Fisher model

Here, we analyse the adaptive dynamics of dispersal and benevolence for the baseline Wright-Fisher model set out in the main text, in which generations do not overlap. Throughout the supporting information, references to equations without the prefix "S" refer to equations of the main text.

#### S.1.1 Directional selection

##### Directional selection on dispersal

We first look at the selection gradient on dispersal. Substituting eqs. (19) and eq. (21) into eq. (6) (with *u = d*) gives the selection gradient on dispersal,

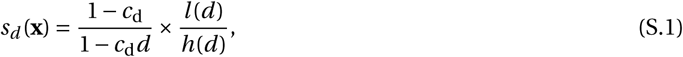

Where

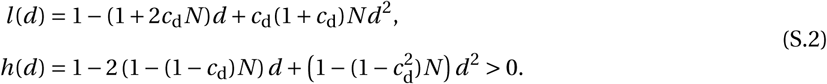

The equilibrium *d*^*∗*^ for dispersal then solves *l* (*d*^*∗*^) *=* 0, which gives

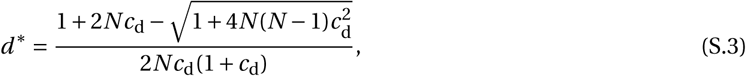

in agreement with previous works on the evolution of dispersal ^e.g., eq. (14) of ref. 54^. In particular, when there is only one adult per site, *N =* 1, the equilibrium eq. (S.3) matches the one found in the classical Hamilton and May model ^5^: *d*^*∗*^ *=* 1/(2 – *p*), where *p =* 1 – *c*_d_ is the probability of survival during dispersal ^eq. (1) of ref. 5^.

The equilibrium eq. (S.3) is an attractor when dispersal is evolving in isolation from *z* if

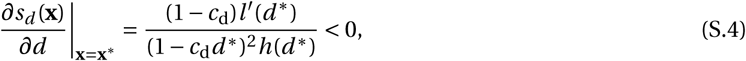

which is always the case because the derivative 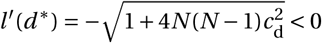 is always negative.

**Directional selection on benevolence.** Substituting eqs. (19) and eq. (21) into eq. (6) (with *u = z*) gives the selection gradient on benevolence,

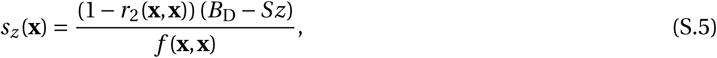

so that the equilibrium **x**^*∗*^ which solves *s*_*z*_ (**x**^*∗*^) *=* 0 is

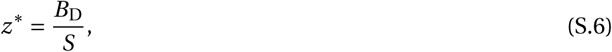

in agreement with previous works on the evolution of *z* alone ^eq. (9) of ref. 23 with κ *=* 0^. Eq. (S.5) shows that dispersal has no effect on the convergence of *z* to an equilibrium here. This is because how limited dispersal influences convergence of *z* depends on the effects of dispersal on genetic relatedness, which favours benevolent behaviour, and on kin competition, which favours self-serving behaviour ^73^, and when local population size is constant (fixed *N*) and generations do not overlap, these effects cancel out regardless of payoffs ^74^.

The equilibrium eq. (S.6) is an attractor point when *z* is evolving alone if

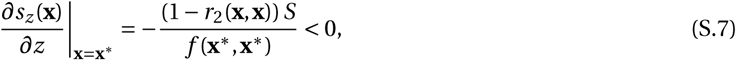

which occurs when *S >* 0 (since 0 *≤ r*_^2^_(**x**, **x**) *<* 1 and *f* (**x**^*∗*^, **x**^*∗*^) *>* 0). So, in order for an internal equilibria to be an attractor, we must have 0 *≤ B*_D_/*S ≤* 1 and *S >* 0, which we will assume for the rest of this section.

**Directional selection when dispersal and benevolence co-evolve** When both traits co-evolve, whether the point **x**^*∗*^ *=* (*z*^*∗*^, *d*^*∗*^) is an attractor depends on the Jacobian matrix (**J**(**x**^*∗*^), eq. 8). Since we find that both offdiagonal entries of the Jacobian are zero, i.e.,

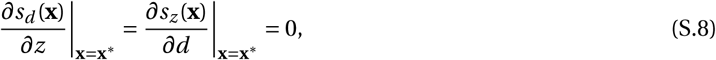

the eigenvalues of the Jacobian **J**(**x**^*∗*^) and of its symmetric part (**J**(**x**^*∗*^)*+***J**(**x**^*∗*^)^T^)/2 are equal to the diagonal entries (eqs. S.4 and S.7). So the conditions for

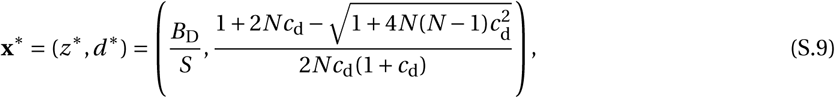

to be an attractor are the same as for *z*^*∗*^ and *d*^*∗*^ to be attractors when evolving in isolation from one another, which we have already derived. Thus, provided 0 *≤ B*_D_/*S ≤* 1 and *S >* 0, the population will converge to **x**^*∗*^ *=* (*z*^*∗*^, *d*^*∗*^) under directional selection.

#### S.1.2 Stabilising selection on dispersal and benevolence when they evolve in isolation from one another

Substituting eqs. (18), (19), (21) and (22) into eqs. (11)-(15) (with *u = v = d*), we find that we can express the quadratic coefficient of selection on dispersal at the equilibrium as a product

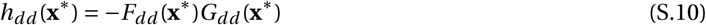

Where

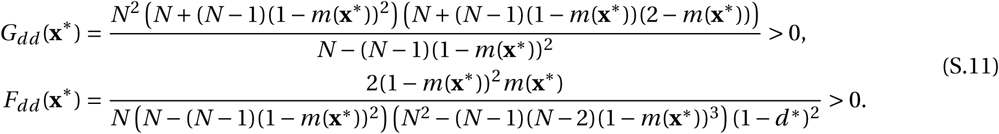

So, *h*_*dd*_ (**x**^*∗*^) *<* 0 is negative, and selection on dispersal when it is evolving alone is always stabilising (as previously found for this life-cycle ^54^).

Similarly, substituting eqs. (18), (19), (21) and (22) into eqs. (11)-(15) (with *u = v = z*), the quadratic coefficient of selection on *z* at the equilibrium can be expressed as

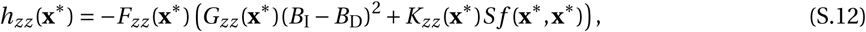

Where

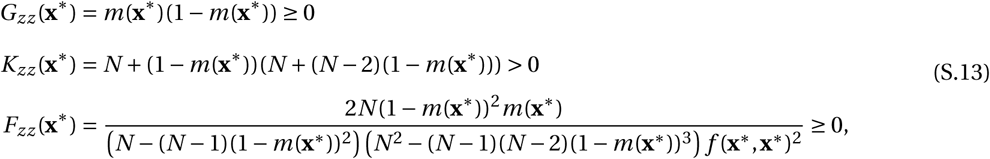

Then, since *S >* 0, *h*_*zz*_ (**x**^*∗*^) *<* 0 is non-positive, and mutations that only change *z* will eventually vanish. So, selection on *z* when it is evolving alone is stabilising for all model parameters.

#### S.1.3 Disruptive selection on dispersal and benevolence when they co-evolve

##### Correlational coefficient of selection

When dispersal and benevolence co-evolve, selection at the equilibrium also depends on the correlational coefficient of selection, and by substituting eqs. (18), (19), (21) and (22) into eqs. (11)-(15) (with *u = z* and *v = d*), we find that it can be expressed as

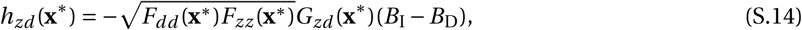

Where

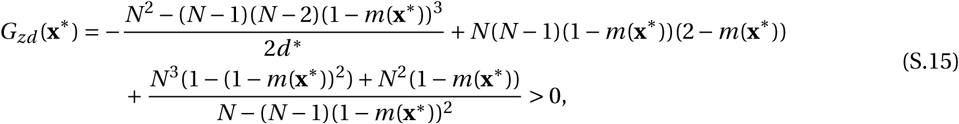

is positive (showing that *G*_*zd*_ (**x**^*∗*^) *>* 0 can be easily achieved with an algebraic computer program ^75^ once we have substituted for the dispersal equilibrium eq. S.3). Since we assume *B*_I_ – *B*_D_ *>* 0, eq. (S.14) shows that the correlational coefficient of selection is negative, *h*_*zd*_ (**x**^*∗*^) *<* 0, which means that selection favours a negative correlation among benevolence and dispersal.

##### Disruptive selection

Substituting eqs. (S.10), (S.12) and (S.14) into eq. (16) shows that the selection will be disruptive whenever

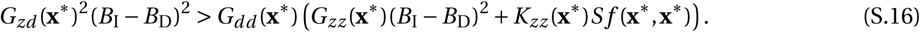

This can be re-arranged to

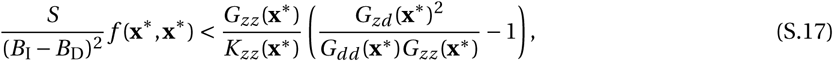

so that the the left hand-side only depends on the properties of the game, and the right hand side only depends on ecological variables: group size *N* and dispersal cost *c*_d_ (with eqs. S.11, S.13, S.15 and eq. 20). Unfortunately, the right hand side of eq. (S.17) is too complicated to be mathematically tractable, but can be analysed numerically (Supplementary Figure S3a). Parameter values that satisfy eq. (S.17) are shown in Figure 2 with *f* (**x**^*∗*^, **x**^*∗*^) *=* 1.

##### Why is selection disruptive?

The above analysis shows that the correlational coefficient of selection can cause polymorphism when *z* and dispersal co-evolve. In order to gain more understanding about this result, let us consider a mutation that arises in a monomorphic population for the equilibrium **x**^*∗*^ and that changes dispersal by Δ_*d*_ *= d*_m_ – *d*^*∗*^ and *z* by Δ_*z*_ *= z*_m_ – *z*^*∗*^. This mutation will invade whenever its growth rate is greater than one, i.e., whenever

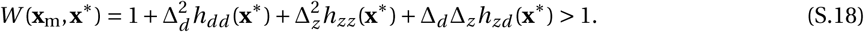

From eq. (S.14), the correlational coefficient of selection *h*_*zd*_ (**x**^*∗*^) *<* 0 is negative. Then, since *h*_*dd*_ (**x**^*∗*^) *<* 0 and *h*_*zz*_ (**x**^*∗*^) *<* 0, eq. (S.18) shows that the only mutations that have a chance of invading are those that have opposite effects on dispersal and *z*, i.e., mutations with Δ_*d*_ Δ_*z*_ *<* 0. Thus, two types of mutation can invade the population, one type that increases *z* and decreases *d* and another type that decreases *z* and increases *d*. Further insights can be brought by looking at the correlational coefficient of selection when dispersal cost is low (i.e., when *c*_d_ is small),

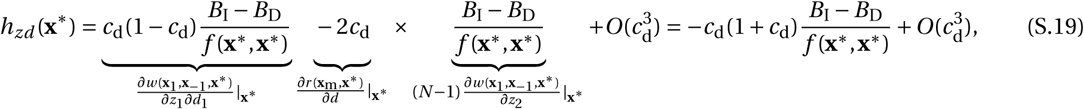

which reveals that what causes the correlational coefficient of selection to be negative, is the combination of the effect of dispersal on pairwise relatedness (*∂r* (**x**_m_, **x**^*∗*^)/*∂d*) with the indirect fitness effects of social interactions (*∂w* (**x**_^1^_, **x**_–1_, **x**^*∗*^)/*∂z*_^2^_), which are proportional to the relative effect of benevolence *B*_I_ – *B*_D_. The correlational coefficient of selection therefore favours mutations or morphs that ensure that its carriers either provide (1) more indirect benefits to their own kin; or (2) less indirect benefits to non-kin.

### S.2 Adaptive dynamics under the Moran model of overlapping generations

Here, we analyse the adaptive dynamics of dispersal and benevolence under the Moran model in which a single individual is replaced in each group at each generation, so that generations overlap.

#### S.2.1 Directional selection

##### Directional selection on dispersal

Substituting eq. (24) and (25) into eq. (6) (with *u = d*) yields the selection gradient on dispersal,

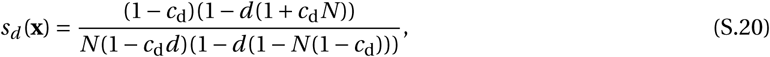

which shows that the equilibrium probability of dispersal (*d*^*∗*^ such that *s*_*d*_ (*d*^*∗*^) *=* 0) is

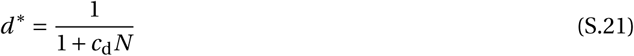

(in agreement with previous work ^eq. (26) of ref. 16^), which is greater than the equilibrium when generations do not overlap (eq. S.3) because kin competition is greater when generations overlap ^16^.

If only dispersal is evolving, the equilibrium eq. (S.21) is a local attractor when

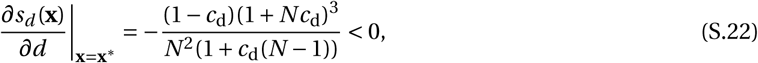

which is always true.

##### Directional selection on benevolence

Substituting eq. (24) and (25) into eq. (6) (with *u = z*), and evaluated at dispersal equilibrium *d*^*∗*^ *=* 1/(1 *+ c*_d_ *N*) gives the selection gradient on the probability *z* of adopting benevolent behaviour B:

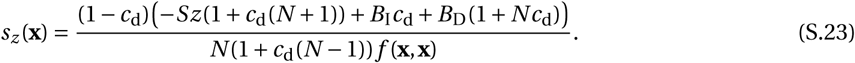

Solving the above for zero gives the equilibrium

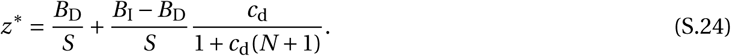

Unlike when generations do not overlap (eq. S.6), group size *N* and dispersal cost *c*_d_ now affect the equilibrium benevolence. This is because these variables determine dispersal (eq. S.3), and when generations overlap, dispersal has greater effects on genetic relatedness than on kin competition ^76^. As a consequence, the indirect fitness benefits of *z*, as measured by *B*_I_ – *B*_D_ *>* 0, play a larger role on its evolution, and the equilibrium benev-olence *z*^*∗*^ is greater when generations overlap than when they do not (eq. S.24 versus eq. S.6).

When only benevolence is evolving, equilibrium eq. (S.24) is a local attractor if

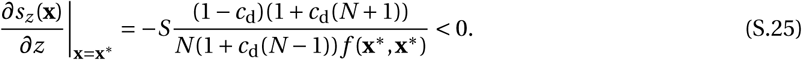

Therefore, the equilibrium for benevolence is internal (0 *< z*^*∗*^ *<* 1) and an attractor when only benevolence is evolving if

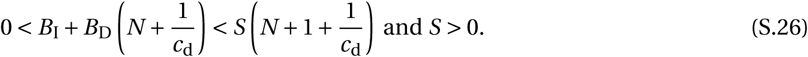

Thus, unlike when generations do not overlap, a cooperative behaviour such that *B*_D_ *<* 0 (like cooperate in the Prisoner’s dilemma game) here can have a convergence stable internal equilibrium.

##### Directional selection when dispersal and benevolence co-evolve

When dispersal and benevolence coevolve, whether the equilibrium

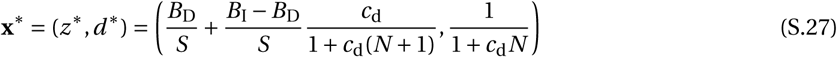

is an attractor depends on the Jacobian matrix **J**(**x**^*∗*^) (eq. 8). We find that the joint equilibrium eq. (S.27) is convergence stable if in addition to eq. (S.26),

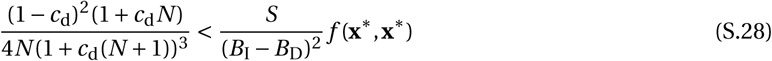

holds, which we will henceforth assume.

#### S.2.2 Stabilising selection on dispersal and benevolence when they evolve in isolation from one another

Once the population has converged to the equilibrium eq. (S.27), the quadratic coefficient of selection on dispersal is found by substituting eqs. (23)-(26) into the Hessian entry eqs. (11)-(15) (with *u = v = d*), which we can express as

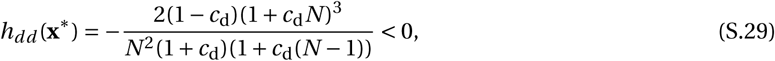

which is always negative. Selection on dispersal alone is therefore stabilising for all model parameters.

Similarly, the quadratic coefficient of selection on benevolence at the equilibrium eq. (S.27) is found by substituting eqs. (23)-(26) into the Hessian entry eqs. (11)-(15) (with *u = v = z*), and we find it can be written as

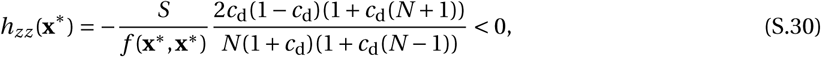

which is always negative since *S >* 0 at an internal attractor equilibrium (eq. S.26). Selection on *z* alone is thus also stabilising for all model parameters.

#### S.2.3 Disruptive selection on dispersal and benevolence when they co-evolve

When dispersal and benevolence co-evolve, whether selection is disruptive depends on the correlational coefficient of selection, which is found by substituting eqs. (23)-(26) into the Hessian entry eqs. (11)-(15) (with *u = z* and *v = d*), and which can be expressed as

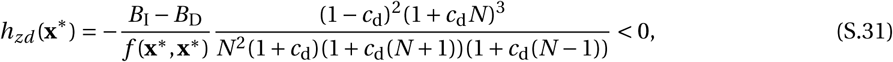

which is always negative when *B*_I_ – *B*_D_ *>* 0. Hence, as in the Wright-Fisher model, correlational selection always favours a negative correlation among benevolence and dispersal in the Moran model.

Substituting eqs. (S.29), (S.30) and (S.31) into eq. (16) reveals after some re-arrangements that whenever

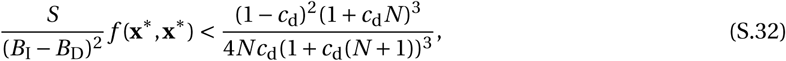

selection is disruptive. Unlike the right-hand side of eq. (S.17), the right-hand side of eq. (S.32) is always positive (since *c*_d_ *<* 1, see Supplementary Figure S3). This means that unlike when generations do not overlap, for any ecological parameters *N* and *c*_d_, there exists a game that leads to disruptive selection when generations overlap. In addition, since cooperative games with *B*_D_ *<* 0 (like the Prisoner’s dilemma game) can have an internal convergence stable equilibrium when generations overlap (eq. S.27), it is possible for such games to lead to polymorphism when generations overlap, which was not the case in our baseline model.

#### S.2.4 Selection when dispersal is cost-free

When dispersal is cost-free (*c*_d_ *=* 0), analysis is much simplified. The population first converges to the equilibrium point

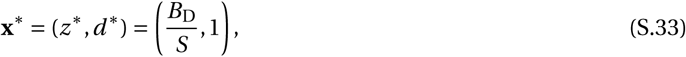

at which there is no kin structure since dispersal is full (*d*^*∗*^ *=* 1). In other words, the population is well-mixed and individuals interact at random. At this equilibrium, the Hessian matrix simply reads as,

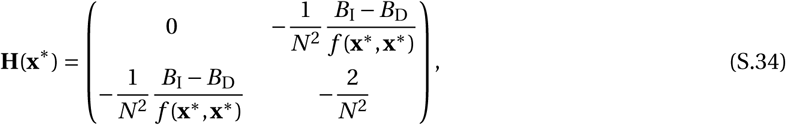

and its dominant eigenvalue is,

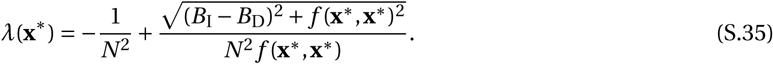

Then, since

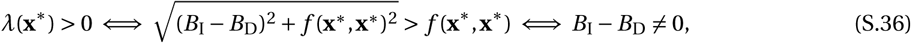

disruptive selection always occurs and leads to a polymorphism that creates kin structure: benevolent individuals that have *z* greater than *z*^*∗*^ *= B*_D_/*S* also disperse less, allowing for relatedness among benevolent individuals to build up. By contrast, when dispersal is fixed at one (*d =* 1), the population will remain monomorphic for the convergence stable equilibrium *z*^*∗*^ for Matrix games ^77^.

### S.3 Individual based simulations

#### Simulations for the baseline model

We performed individual based simulations for a population composed of *N*_d_ *=* 1250 groups, each populated by *N =* 8 haploid individuals, using Mathematica 11.0.1.0^75^. Starting with a population monomorphic for no dispersal *d =* 0 and full benevolence *z =* 1, we track the evolution of the multidimensional phenotypic distribution under the constant but small influx of mutations. Each individual *i =* 1,…, 10000 is characterised by a dispersal value *d*_*i*_ and a benevolence value *z*_*i*_. At the beginning of a generation, we calculate the fecundity *f*_*i*_ of each individual according to its benevolence and that of its neighbours (eq. 17, with *f*_0_ *=* 1). Then, we form the next generation of adults by sampling *N* individuals in each group with replacement, where each individual from the parental generation is weighted according to whether they belong to the group on which the breeding spot is filled or not. If an individual belongs to the same group in which a breeding spot is filled, then its weight is *f*_*i*_ (1 – *d*_*i*_). If it belongs to another group, then its weight is *f*_*i*_ *d*_*i*_ (1 – *c*_d_)/(*N*_d_ - 1). Once an individual is chosen to fill the breeding spot, each locus independently mutates with probability 0.01. If they do not mutate, then the offspring has the same phenotypic values as its parent. If they mutate, then we add to parental values a small perturbation that is sampled from a normal distribution with mean 0 and variance 0.01^2^. The resulting phenotypic values are truncated to remain between 0 and 1. We repeated the procedure for 3 *×* 10^4^ generations.

#### Fixed recombination

We incorporated recombination in our model by adding a stage of diploidy in the lifecycle of our population. After social interactions within groups, haploid adult individuals produce haploid gametes that disperse according to their genotype at the dispersal locus. Then, gametes randomly fuse within groups to make a diploid zygote who undergoes meiosis and whose genome recombine according to a probability *R*. A diploid zygote then produces two haploid individuals one of which (randomly sampled) develops into an adult. In terms of simulations, this modified life-cycle entails that after calculating the fecundity *f*_*i*_ of each individual, each of the *N* breeding spots are filled by first sampling two haploid gametes from the parental generation according to the weighted sampling described in **Simulations for baseline model**, and fuse to make a diploid zygote. Then, say a diploid zygote has parents indexed *i* and *j* with genotypes (*z*_*i*_, *d*_*i*_) and (*z*_*j*_, *d*_*j*_) respectively. With a probability *R*, recombination occurs in this zygote, which produces two recombinant haploid gametes, (*z*_*i*_, *d*_*j*_) and (*z*_*j*_, *d*_*i*_). With complementary probability 1*-R*, recombination does not occur and the two haploid gametes are the parental ones, (*z*_*i*_, *d*_*i*_) and (*z*_*j*_, *d*_*j*_). The breeding spot is filled by sampling one of the haploid gamete at random (with equal probability) which develops into an adult individual. The rest of the simulation is as in **Simulations for baseline model**.

#### Recombination evolution

In order to model recombination evolution, we added a third quantitative locus that controls the recombination probability at the diploid stage. The genome of a haploid individual therefore carries three quantitative loci: (*z*, *d*, *τ*), where as before *z* encodes the probability of adopting B action, *d* the probability of dispersal at birth, and *τ* controls the probability of recombination among *z* and *d* loci. For simplicity, we assume that the *τ* locus is completely linked to the *d* locus. Two alleles segregate at the recombination locus *τ*: a wild-type *R* which codes for a recombination probability of 1/2 and a dominant mutant *r* for zero recombination. Diploid zygotes are formed as above. Then, say a diploid zygote has parents indexed *i* and *j* with genotypes (*z*_*i*_, *d*_*i*_, *τ*_*i*_) and (*z*_*j*_, *d*_*j*_, *τ*_*j*_) respectively. Recombination in this zygote occurs with a probability 1/2 if *τ*_*i*_ *= τ*_*j*_ *= R*, or zero otherwise. If recombination occurs, the haploids produced are (*z*_*i*_, *d*_*j*_, *τ*_*j*_) and (*z*_*j*_, *d*_*i*_, *τ*_*i*_) since the *τ* locus is completely linked to the *d* locus. Mutation occurs at each locus independently with probability 0.01. Quantitative effects of mutation at the *z* and *d* loci are as in **Computer simulations for baseline model**, while mutations at the *τ* locus changes *r* to *R* and *R* to *r*. The rest of the simulation is as in **Computer simulations for baseline model**.

*F*_ST_ **calculations.** In order to calculate the *F*_ST_ measures of genetic differentiation shown in Figure 5 at a given generation, we first calculated the average probability of adopting the benevolent action in the population at that generation,

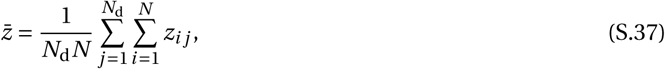

where *N*_d_ is the number of groups in the population and *z*_*i j*_ is the average genotypic value at the *z* locus (i.e., the probability of adopting the benevolent action) of individual indexed *i ∈* {1,…, *N*} in group indexed *j ∈* {1,…, *N*_d_}. Next, we divided the population between a benevolent and self-serving morph by comparing each individual to the population average 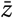. Those that carried a genotypic value *z* greater than 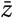 were considered to be from the benevolent morph, and those with a genotypic value less than 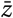, from the self-serving morph. The set of individuals belonging to the benevolent morph in a group *j ∈* {1,…, *N*_d_} is then described by the set

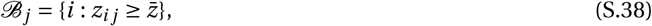

which gives the individual indices *i* of all the individuals from the benevolent morph in group *j*. Similarly, the set of individuals belonging to the self-serving morph in a group *j ∈* {1,…, *N*_d_} is

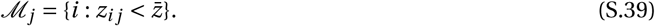

We denote by *|*ℬ_*j*_ *|* the size of set ℬ _*j*_, i.e., the number of individuals from the benevolent morph in group *j*, and similarly, by *|ℳ*_*j*_ *|*, the number from the self-serving morph in group *j*.

Let us first consider the procedure to calculate the genetic differentiation among morphs at the selected locus, i.e., at the *z* locus (full black curve in Figure 5). We first calculated the genetic variance among morphs as

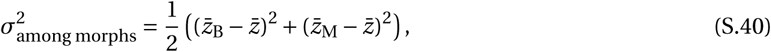

Where

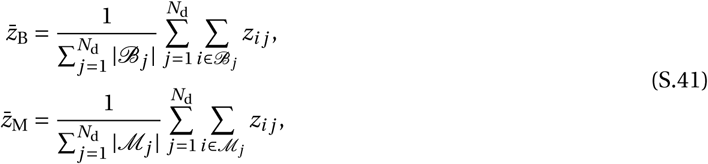

respectively give the average *z* genotypic value within the benevolent and self-serving morph. Next, we calculated the genetic variance within morphs as

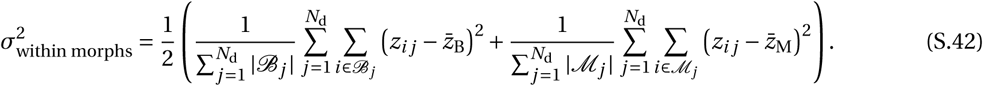

The genetic differentiation among morphs at the selected locus is then

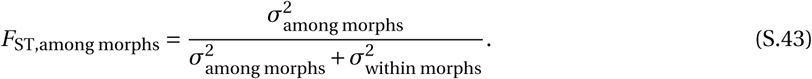

We also calculated the genetic differentiation among groups, within each morph (full blue and red curves in Figure 5). We first calculated the genetic variance among groups, within the benevolent morph, as

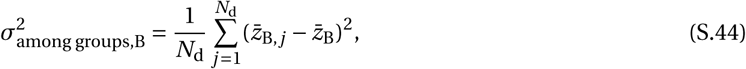

Where

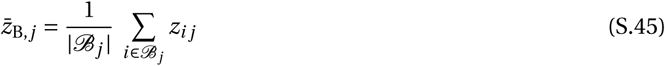

is the average *z* genotypic value in the benevolent morph in group *j*. Second, we calculated the genetic variance within groups, within the benevolent morph, as

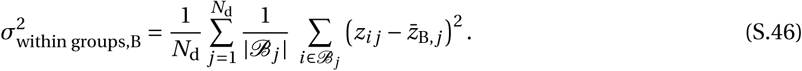

The genetic differentiation among groups, within the benevolent morph, at the selected locus (full blue curve in Figure 5) is then given by

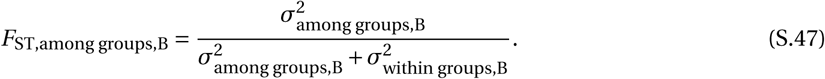

Similarly, genetic differentiation among groups, within the self-serving morph, at the selected locus (full red curve in Figure 5) is given by

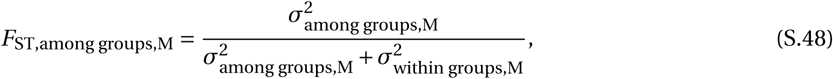

Where

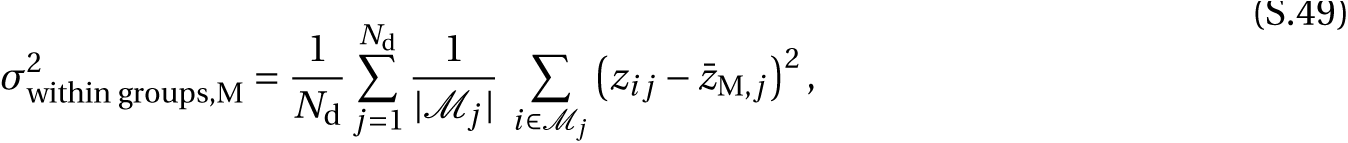

and

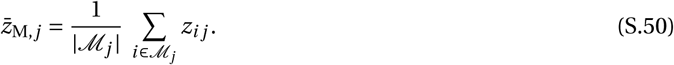

Measures of genetic differentiation at the neutral locus (dashed curves in Figure 5) are calculated the same way as above (eqs. S.40-S.50), replacing the genotypic values at the *z* locus by those at the neutral locus.

